# Remotely sensed carotenoid dynamics predict photosynthetic phenology in conifer and deciduous forests

**DOI:** 10.1101/2021.04.10.439296

**Authors:** Christopher Y.S. Wong, Lina M. Mercado, M. Altaf Arain, Ingo Ensminger

## Abstract

Crucially, the phenology of photosynthesis conveys the length of the growing season. Assessing the timing of photosynthetic phenology is key for terrestrial ecosystem models for constraining annual carbon uptake. However, model representation of photosynthetic phenology remains a major limitation. Recent advances in remote sensing allow detecting changes of foliar pigment composition that regulate photosynthetic activity. We used foliar pigments changes as proxies for light-use-efficiency (LUE) to model gross primary productivity (GPP) from remote sensing data. We evaluated the performance of LUE-models with GPP from eddy covariance and against MODerate Resolution Imaging Spectroradiometer (MODIS) GPP, a conventional LUE model, and a process-based dynamic global vegetation model at an evergreen needleleaf and a deciduous broadleaf forest. Overall, the LUE-models using foliar pigment information best captured the start and end of season, demonstrating that using regulatory carotenoids and photosynthetic efficiency in LUE models can improve remote monitoring of the phenology of forest vegetation.

## Introduction

Assessing global terrestrial photosynthesis or gross primary productivity (GPP) is important for understanding the global carbon cycle. The magnitude of total annual GPP depends strongly on timing: on the onset of photosynthetic activity in spring and its cessation in the autumn ^1^. Therefore, the ability to accurately assess the seasonal dynamics of photosynthesis, or photosynthetic phenology, is key for terrestrial ecosystem models (TEMs) to reduce bias in representing the seasonal cycle of carbon uptake ^2^. Yet how models represent photosynthetic phenology remains uncertain. This uncertainty is due to the lack of a mechanistic understanding of the controls of vegetation phenology, which limits current model algorithms and parameterization ^3,4^. Warming temperatures are also likely to shift the timing of photosynthetic phenology and further impact the performance of TEMs ^5^.

TEMs conventionally model phenology by employing process-based or remote sensing-based approaches. Process-based models, or dynamic global vegetation models (DGVMs), generally represent photosynthesis based on the Farquhar, von Caemmerer and Berry biochemical model of leaf photosynthesis ^6^. An example is the widely used JULES model, where the phenology of photosynthesis in deciduous vegetation is estimated based on the presence or absence of leaves, as a simple proxy for leaf phenology, and on photosynthetic potential ^7^. Both parameters are temperature dependent and thus vary with season, e.g. leaf phenology is derived from accumulated growing degree days above a temperature threshold ^3^. In contrast, evergreen vegetation retains its leaves and hence largely lacks easily discernable seasonal leaf phenology. As a consequence, estimates of photosynthetic phenology simply rely on meteorological factors. Regardless of the type of vegetation, the meteorological constraints of both deciduous and evergreen leaf phenology are complex and difficult to model ^3,8^. Although temperature and photoperiod are generally the major environmental factors controlling leaf phenology, species-specific responses and other environmental factors such as soil moisture and precipitation are also known to contribute to the timing of phenology ^4^.

Remote sensing approaches for assessing phenology offer a way to use direct large-scale observations of canopy structure and vegetation greenness. In many ecosystems, remotely sensed vegetation greenness effectively represents the phenology of photosynthesis ^9^. However, monitoring phenology in evergreen forests remains a challenge as they show little seasonal variation in vegetation greenness, which is often decoupled from the phenology of photosynthesis ^10,11^. Instead, the phenology of photosynthesis in evergreen vegetation is largely driven by invisible physiological adjustments regulating photosynthetic activity ^12–14^. To represent the regulation of photosynthesis, remote sensing-based models integrate both remote sensing and meteorological inputs to estimate GPP.

Models using remote sensing are generally based on the light-use efficiency (LUE) model (Eqn 1) ^15^. In this model, the remotely sensed normalized difference vegetation index (NDVI) is often used to parameterize the fraction of absorbed photosynthetically active radiation (*f*_APAR_). Photosynthetically active radiation (PAR) represents incident radiation while light-use efficiency (ε) is typically estimated from temperature and vapour pressure deficit (VPD) limits ^16^.

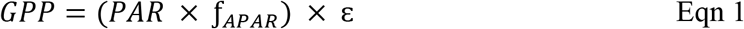

The LUE model shown in Eqn 1 gives an accurate representation of GPP for deciduous vegetation because *f*_APAR_ reflects closely leaf area index (LAI), canopy greenness and photosynthetic activity ^17^. However, evergreen vegetation remains green year-round, and LAI and canopy greenness remain high even when photosynthesis is downregulated during the winter. Therefore, it is important to consider ε as the key indicator representing regulation of photosynthetic activity in evergreen vegetation. Assessing ε remotely has been challenging as it represents the fraction of absorbed PAR that is used in all of the photosynthetic processes ^18,19^. Instead, ε is often represented indirectly, as a fixed biome specific constant ^20^, or directly, as a dynamic ε that varies according to meteorological data, reflecting the effects of temperature and water limitations on photosynthetic activity ^16,21^. The use of meteorological data for parameterizing ε in the LUE model does not fully capture the local heterogeneity of ε and often limits the model’s accuracy ^22^. Therefore, to improve the LUE model, studies have sought to remotely assess ε directly from vegetation ^19,23^.

The regulation of photosynthetic activity and therefore ε is linked to carotenoid pigments involved in the safe dissipation of excess light energy ^24^. Over short time periods - seconds to minutes - photosynthetic activity is regulated through the reversible conversion of the xanthophyll cycle pigment violaxanthin to the photoprotective antheraxanthin and zeaxanthin ^24^. The interconversion of the xanthophyll cycle pigments can be remotely assessed by the photochemical reflectance index (PRI) ^25^. However, over seasonal time periods, PRI predominantly reflects changes in the ratio of carotenoids per chlorophylls (Car/Chl) ^26–28^. Thanks to recent advances in remote sensing, we can detect seasonal Car/Chl using MODIS (moderate resolution imaging spectroradiometer) through an analog vegetation index of PRI, the chlorophyll/carotenoid index (CCI) ^29^. Long-term variation of Car/Chl reflects the physiological effort of vegetation to balance light absorption and photoprotection, which in turn leads to changes in photosynthetic activity. A relative increase in carotenoids corresponds to a decrease in photosynthetic activity ^14,30^ and contributes to sustained dissipation of excess light when photosynthesis is downregulated, a photoprotective mechanism crucially important for overwintering conifers ^12,14,24^. Studies have confirmed that PRI and CCI closely track seasonal photosynthetic activity of both evergreen and deciduous trees ^31–33^. These findings suggest that assessing carotenoid pigment composition remotely through spectral reflectance measurements can be used as a reliable indicator of ε and photosynthetic phenology for northern forests.

Besides the visible reflectance signal used to estimate PRI and CCI, there are also promising approaches using the near-infrared reflectance of vegetation index (NIR_v_) to assess GPP remotely ^34,35^. Since NDVI and NIR_v_ share some reflectance bands, and because the latter relies on near-infrared reflectance to better account for leaf area and canopy structure, NIR_v_ may actually represent the light capture of vegetation and thus the absorbed photosynthetically active radiation APAR. Therefore, the potential role of NIR_v_ as a substitute for NDVI in the LUE model should be evaluated.

This study aims to use carotenoid-based vegetation indices to improve the prediction of photosynthetic phenology and to increase the accuracy of GPP estimates in conifer and deciduous forests. For this purpose, we modified the conventional LUE using the vegetation indices NDVI and NIRV_v_ as proxies for light harvesting and PRI and CCI as proxies for photosynthetic efficiency; tower-based eddy covariance and satellite-based remote sensing data were used to evaluate a temperate evergreen needleleaf forest (ENF) and a deciduous broadleaf forest (DBF). The accuracy of the modified LUE models was validated by comparing GPP derived from our and from conventional models with the observed GPP obtained from eddy covariance measurements at the ENF and DBF sites.

## Results

### Comparing observed GPP with modelled GPP

We compared stand level photosynthesis expressed as observed gross primary productivity (GPP) from eddy covariance measurements with modelled GPP from the period 2012 to 2019 (where data were available) in a temperate evergreen needle forest (ENF) and a temperate deciduous broadleaf forest (DBF) (Fig. 1). Overall, the seasonal pattern of GPP reveals that the ENF had a longer growing season (March/April to late December) compared to the DBF (May to late October), leading to total annual GPP that was about 20% larger in the ENF than in the DBF (Fig. 2). Notably, estimates of total annual GPP derived from vegetation index-based LUE models provided a better approximation of mean total annual GPP_obs_ than MODIS_GPP_, LUE_Met_ and JULES_Farq_ at both sites during the period 2015 to 2018 at both forest sites (Fig. 2).

**Figure 1.**
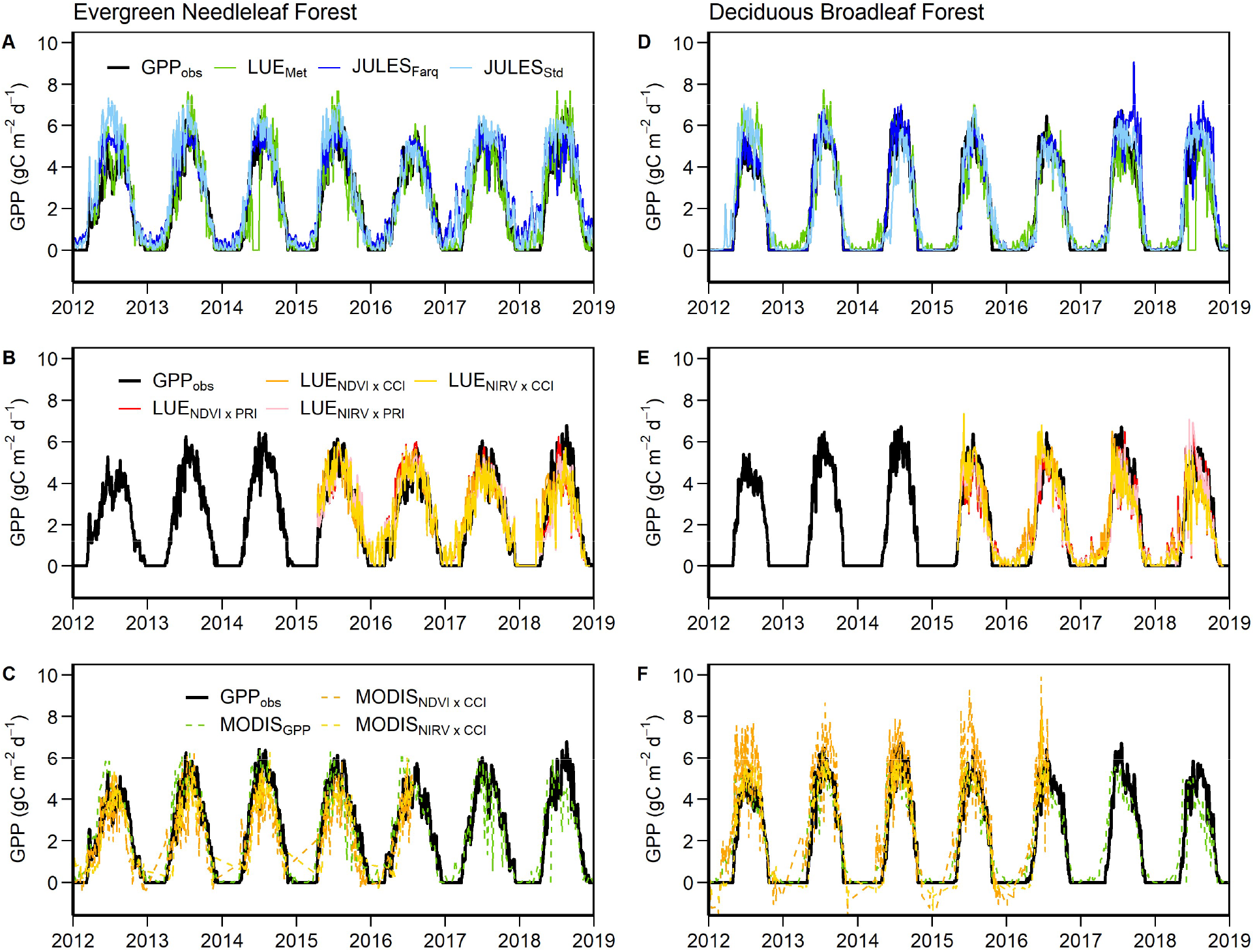
Seasonal pattern of stand level photosynthesis (expressed as daily GPP_obs_, black line) and modelled daily GPP for an evergreen needleleaf forest (A-C) and a deciduous broadleaf forest (D-F) from 2012 to 2019. Models shown include LUE_Met_ (green), JULES_Farq_ (blue), JULES_Std_ (cyan), LUE_NDVI x PRI_ (red), LUE_NDVI x CCI_ (orange), LUE_NIRv x PRI_ (pink), LUE_NIRv x CCI_ (yellow), MODIS_GPP_ (green), MODIS_NDVI x CCI_ (pink), and MODIS_NIRv x CCI_ (yellow). Top row represents meteorological-based models, middle row represents canopy scale vegetation index models, and bottom row represents satellite-based MODIS models. Lines represent the 5 day running means. Model details are provided in Table S3.

**Figure 2.**
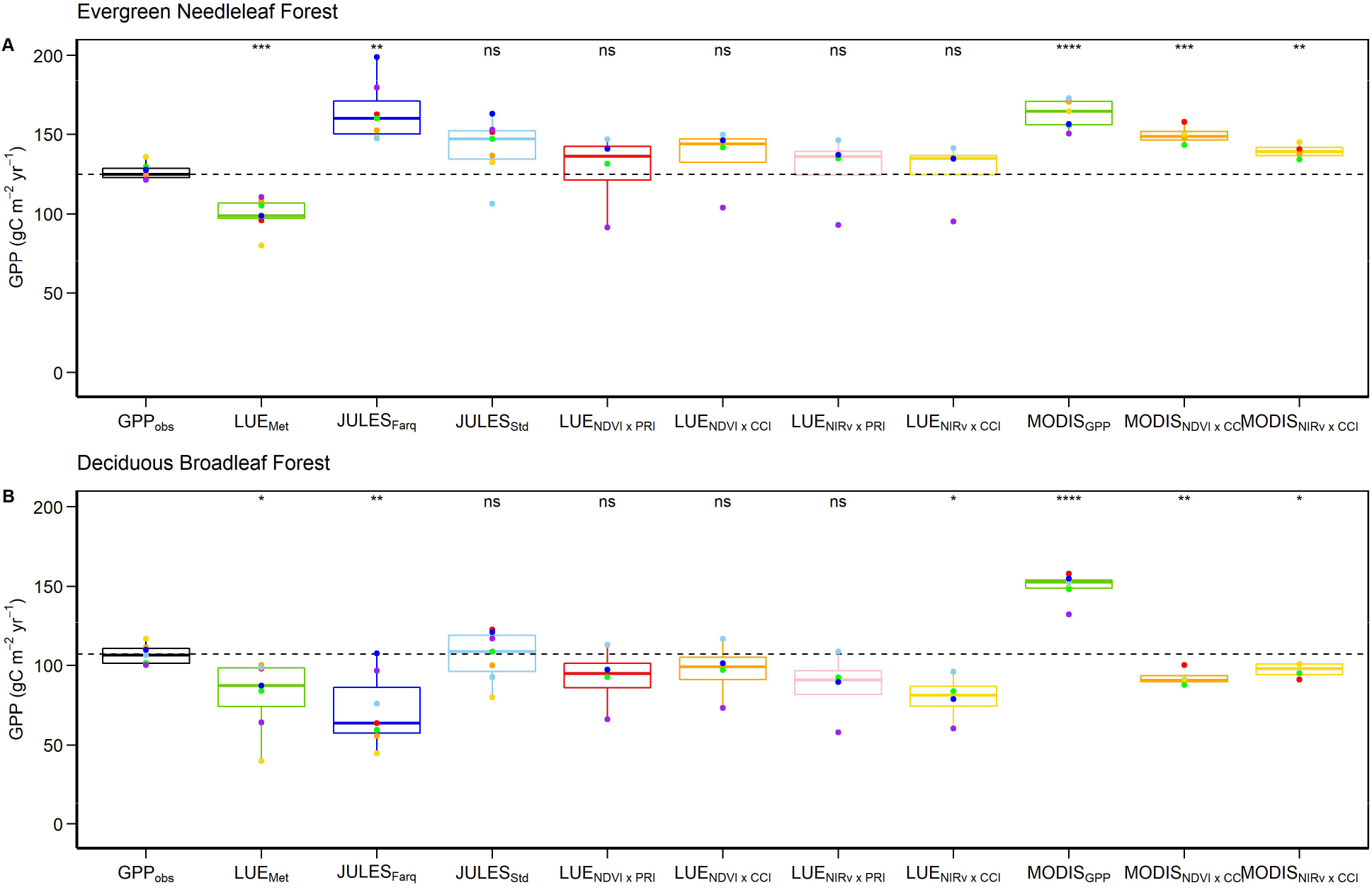
Difference in annual stand level photosynthesis (expressed as sum of daily GPP_obs_) and modelled daily GPP for an evergreen needleleaf forest (A) and a deciduous broadleaf forest (B). Models shown include LUE_Met_ (green), JULES_Farq_ (blue), JULES_Std_ (cyan), LUE_NDVI x PRI_ (red), LUE_NDVI x CCI_ (orange), LUE_NIRv x PRI_ (pink), LUE_NIRv x CCI_ (yellow), MODIS_GPP_ (green), MODIS_NDVI x CCI_ (pink), and MODIS_NIRv x CCI_ (yellow). Boxes display the first quartile, median, and third quartile, whiskers display the range. Dots represent annual GPP for 2015 (green), 2016 (orange), 2017 (blue) and 2018 (red). The median of GPP_obs_ (dashed line) is used as baseline to illustrate the deviation of the models from observed annual GPP. Asterisks denote significant differences between modelled data compared to GPP_obs_ data determined by a Student’s t test. ns = non significant; *P < 0.05; **P < 0.01; ***P < 0.001. Model details are provided in Table S3.

However, comparisons of observed and modelled GPP at daily time scales, demonstrates that all models provided accurate estimates of daily stand level GPP; R^2^ ranged from 0.45 to 0.88, with the highest R^2^ obtained for the JULES_Farq_ model for the ENF (R^2^ = 0.81) and the LUE_Met_ model for the DBF (R^2^ = 0.88) (Fig. 3). For most models, the seasonal variability of daily stand GPP was better represented in the DBF than in the ENF. Although, vegetation index-based LUE models showed slightly lower R^2^ compared to the LUE_Met_ and JULES models (Fig. 3), the vegetation index-based LUE models performed more uniformly between the ENF and DBF with regard to slopes and intercepts. Model performance and their ability to reflect the seasonal variation of stand photosynthesis was evaluated and demonstrates that all models represent the autumn downregulation more accurately than the spring recovery (Table S1). Further analysis of model GPP residuals indicated that the largest bias occurs near the peak of the growing season (Fig. S1) at high temperatures (Fig. S2) and PAR (Fig. S3).

**Figure 3.**
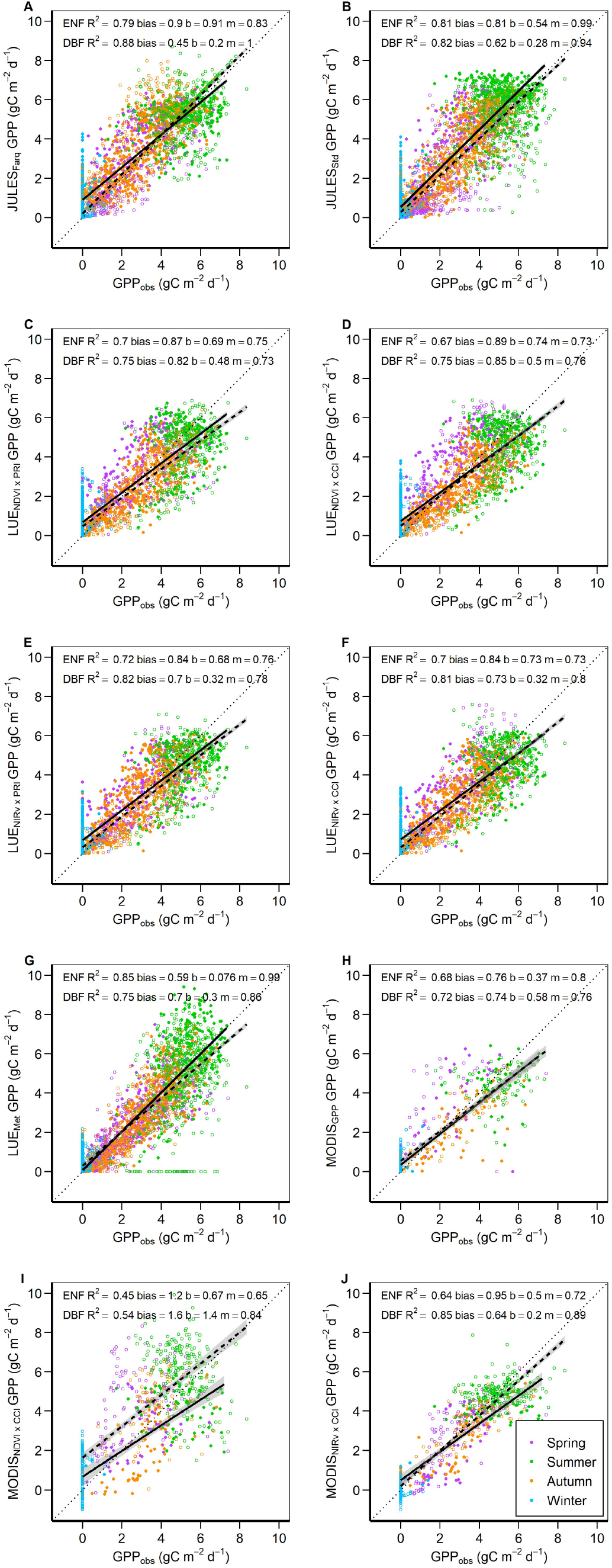
Relationship between observed stand photosynthesis (expressed as daily GPP_obs_) and modelled daily GPP for an evergreen needleleaf forest (ENF; closed symbols and solid line) and a deciduous broadleaf forest (DBF; open symbols and dashed line) from 2012 to 2019. Models shown include JULES_Farq_ (A), JULES_Std_ (B), LUE_NDVI x PRI_ (C), LUE_NDVI x CCI_ (D), LUE_NIRv x PRI_ (E), LUE_NIRv x CCI_ (F), LUE_Met_ (G), MODIS_GPP_ (H), MODIS_NDVI x CCI_ (I), and MODIS_NIRv x CCI_ (J). Colors of datapoints reflect different seasons as determined by the phenology dates of GPP_obs_. Shaded regions indicate 95% confidence interval around the line of best fit. Reported are the coefficient of determination (r^2^), bias, intercept (b) and slope (m) of each line of best fit representing all data. Sample size are: JULES n = 1657 (ENF), n = 2557 (DBF); LUE n = 1093 (ENF), n = 1125 (DBF); MODIS GPP n = 211 (ENF), n = 322 (DBF); and MODIS_NDVI x CCI_ and MODIS_NIRv x CCI_ n = 184 (ENF), n = 518 (DBF). All relationships are statistically significant at P < 0.001. Model details are provided in Table S3.

### Timing of photosynthetic phenology

Using the mathematical derivatives ^36^, we determined the start of growing season (SOS) and its end (EOS), which are key metrices of the phenology of photosynthesis. We observed considerable differences across the models in their ability to match the timing of observed SOS or observed EOS that was determined from the eddy covariance data (GPP_obs_) (Fig. 4, S4) with maximum differences between observed and modelled timing of phenology of up to 40 days (Figs. 4). Importantly, for ENF, the vegetation index-based LUE models produced the most accurate estimates of modelled SOS and EOS, consistently ranging within 10 and 20 days, respectively, of observed SOS and EOS (Fig. 4A,B). The LUE_Met_, JULES and MODIS_GPP_ models generally revealed greater annual variability in the timing of photosynthetic phenology. For example, SOS derived from the JULES models advanced SOS of GPP_obs_ by 20 days (Fig. 4A). Similar to the ENF, the vegetation index-based LUE models provided the most accurate determination of SOS for the DBF, with dates typically being within 10 days of observed SOS and with little variation between years (Fig. 4C). The LUE models predicted the dates for EOS with a delay of about 10 days compared to observed EOS and with little interannual variation (Fig. 4D). In contrast to the vegetation index-based LUE models, the SOS and EOS from the LUE_Met_ revealed much larger deviations from observed SOS and EOS and larger interannual variability. The largest interannual variability was determined for the phenological dates extracted from the JULES models, ranging from −35 to + 25 days for SOS and +10 to +35 days for EOS (Fig. 4C,D). MODIS_GPP_ captured SOS earlier by 30 days and showed an EOS delayed by 15 days with large annual variability (Fig. 4C,D).

**Figure 4.**
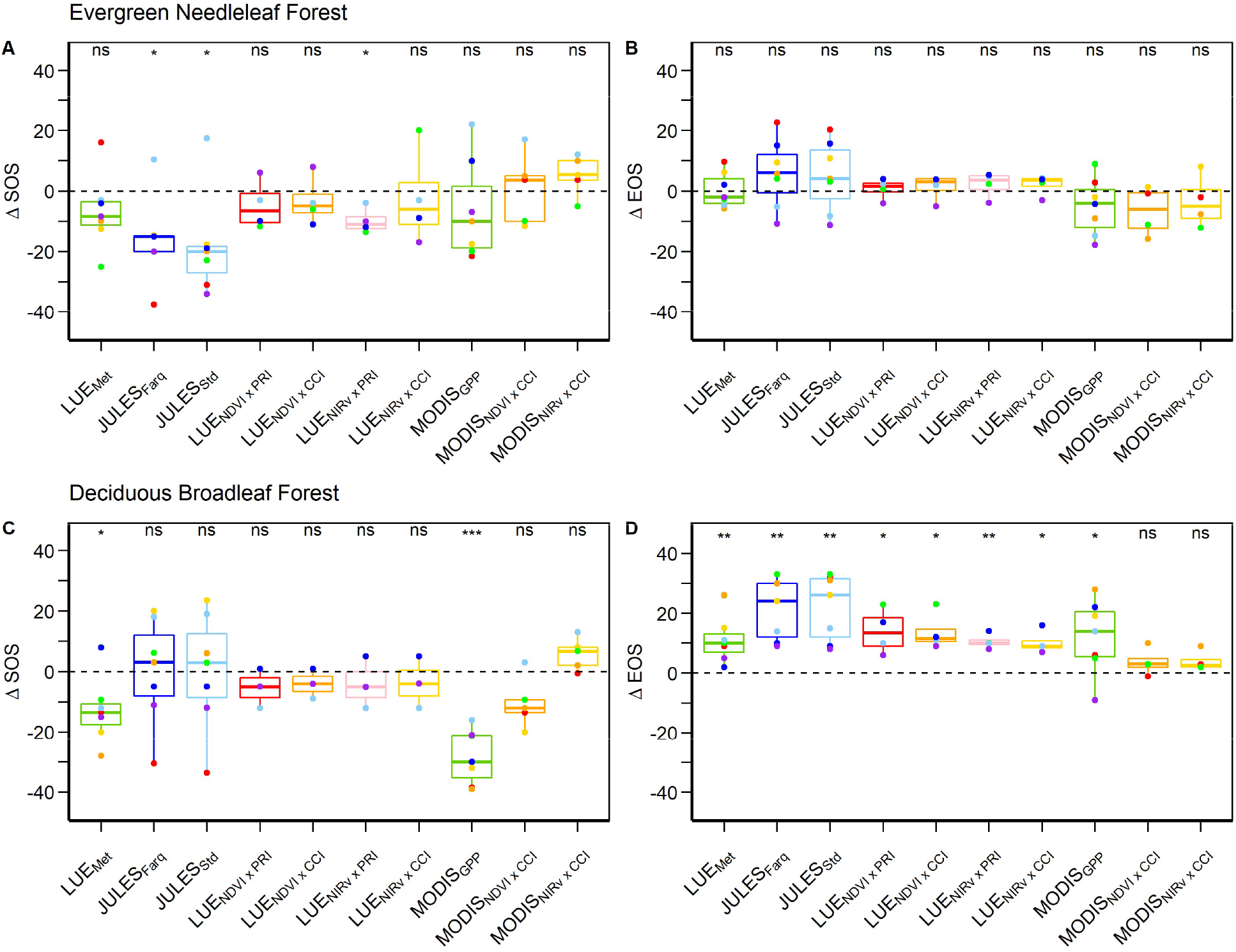
Difference in the timing of photosynthetic phenology in an evergreen needleleaf forest (A,B) and a deciduous broadleaf forest (C,D). Start (Δ SOS) and end (Δ EOS) of the growing season dates determined from 2012 to 2019 observed GPP_obs_ are shown as a reference by the dashed line, boxplots show the deviation of Δ SOS and Δ EOS derived from modelled GPP. Boxes display the first quartile, median, and third quartile, whiskers display the range. Dots represent the difference in number of days for 2012 (red), 2013 (orange), 2014 (yellow), 2015 (green), 2016 (light blue), 2017 (blue) and 2018 (purple). Negative values denote earlier onset of modelled SOS and EOS, positive values denote delay of modelled SOS and EOS compared to observed SOS and EOS from GPP_obs_. Asterisks denote a significant difference between modelled and observed Δ SOS and Δ EOS dates determined by a Student’s t test. ns = non significant; *P < 0.05; **P < 0.01 ***P < 0.001.

## Discussion

How to determine large-scale phenology and accurately predict GPP remains a challenge for TEMs. Generally, TEMs overestimate the growing season length with early SOS and EOS delayed up to two weeks or more ^1,2^. Our study demonstrates that vegetation index-based LUE models with PRI or CCI as proxies for photosynthetic efficiency provide improved objective measures of the phenology of GPP_obs_ in a temperate evergreen needleleaf forest and deciduous broadleaf forest. Conventional process-based and LUE models rely on simplified relationships between meteorological data and photosynthetic activity to constrain photosynthesis in response to environmental conditions. However, these relationships are difficult to model, given the complexity of the spatial, temporal and ecological processes that determine GPP over time ^1,3^. Increasing model complexity may improve model performance as observed in higher R^2^ in the JULES_Farq_ model compared to the JULES_Std_ model (Fig. 3A,B). However, increased complexity may introduce additional model parameterization uncertainty and limit plant functional types that exist outside of model calibration data ^37^.

Compared to the other models, the simple vegetation index-based LUE model performed as well as a complex model in estimating daily stand photosynthesis (Fig. 3), but also revealed improved estimation of SOS and EOS (Fig. 4). In the vegetation index-based LUE model, PRI and CCI are proxies for photosynthetic efficiency, allowing vegetation function to be directly assessed compared to a conventional meteorological representation of photosynthetic efficiency. PRI and CCI are sensitive to the ratio of foliar carotenoid and chlorophyll pigments ^28,29,33^. Carotenoid pigments are involved in the photoprotection of light-harvesting complexes and thus help regulate both short- and long-term photosynthetic activity ^24,30^. Because the role of carotenoid and chlorophyll pigments across evergreen and deciduous trees is conserved, their use as proxies for photosynthetic phenology is robust and accurate ^31,33^.

The light-harvesting component in the LUE model often uses NDVI as a proxy for *f*_APAR_ due to its sensitivity to chlorophyll content and light absorption ^16,17^. Unlike NDVI, the role of NIR_V_ in the LUE model is unclear. Originally, Badgley, et al. ^34,35^ used NIR_V_ as a direct proxy for GPP, one that approximates NIR reflectance to minimize the influence of background signals and the structural effects of the canopy. However, Wang, et al. ^38^ suggest that seasonal NIR_V_ may reflect the greening-up of vegetation and hence the capacity for light harvesting rather than GPP itself. Evaluating the role of NIR_V_ as potential proxy for *f*_APAR_, APAR or GPP, we found that the strongest relationships were between NIR_V_ and APAR (Fig. S5). However, the MODIS NIR_V_ product exhibited a weaker relationship in the evergreen needle forest (Fig. S5H), largely due to a seasonal hysteresis in the relationship between spring recovery (nonlinear) versus autumn downregulation (linear) (Fig. S6). This discrepancy may be due to differences in spring recovery and autumn downregulation for chlorophyll pigment and leaf area index (LAI) ^39^, a conclusion that supports the results of Wang, et al. ^38^ in prairie grasslands. Overall, using NIR_V_ as a proxy for APAR in the LUE model enhances performance (Fig. 3) and has the added benefit of further simplifying the LUE model by reducing the need to include PAR.

The simple LUE models based on a vegetation index improved the estimation of the phenology of photosynthesis in both the ENF and DBF (Fig. 4). This suggests that the use of vegetation indices as proxies for light harvesting (via NDVI or NIR_V_) and photosynthetic efficiency (via PRI or CCI) for the detection of phenology can be used across different types of vegetation and is not restricted to evergreen forests. Since the roles of chlorophyll and carotenoid pigments in photosynthetic activity are conserved across higher plant species and plant functional types, we suspect that PRI or CCI will remain effective proxies for photosynthetic efficiency across plant functional types ^23^. In addition to the phenology of photosynthesis, PRI and CCI may also act as indicators of stress events (e.g. drought), which affect photosynthetic efficiency ^40^. Given the important role of carotenoid pigments in the regulation of photosynthesis across short (diurnal) and long timescales (seasons), the use of PRI/CCI as proxies for ε in the LUE model can be advantageous for reflecting local site-specific photosynthetic activity. Therefore, further evaluation of vegetation index-based LUE models is recommended across both spatial and temporal scales in addition to additional regions and ecosystems. Advances in the reprocessing of MODIS data from the Aqua and Terra satellites using Multi-Angle Implementation of Atmospheric Correction (MAIAC) ^41^, are enabling CCI to be used in further large-scale studies and, potentially, on the global scale.

An alternative satellite-based approach using solar-induced fluorescence (SIF) has also shown promise in assessments of the photosynthetic phenology of northern forests ^10^. However, because SIF is relatively new, its role in the LUE model is unclear. SIF appears to contain information on both absorbed PAR and chlorophyll content as well as ε ^10,42^. SIF and PRI/CCI may represent the distribution of absorbed light energy via chlorophyll fluorescence and non-photochemical quenching, respectively ^43,44^. Currently, SIF and CCI satellite products are hard to compare as they are obtained from different satellite platforms. Combining these datasets will lead to considerable loss of spatial and temporal resolution. Therefore, paired measurements of SIF and vegetation indices have been proposed as part of the Fluorescence Explorer (FLEX) mission expected to launch in 2022 ^45^, which could lead to SIF and PRI/CCI models being explored for their potential to assess the phenology of GPP from space.

Our findings expand on the ability of the recently described CCI ^29^ to act as a proxy for ε and that of the recently described NIR_v_ ^35^ to act as a proxy for APAR in a simple LUE model. Although conventional LUE and process-based models poorly reflect the timing of phenological events, we have shown that a PRI- or CCI-based LUE model may improve how the phenology of photosynthesis is detected. The increasing temperatures resulting from climate change may have implications for the ability of current LUE and process-based models to accurately predict phenology. Temperature and photoperiod are the environmental cues triggering phenology, but climate warming will result in asynchronous phasing of photoperiod and temperature during the spring and autumn ^12^. This asynchrony will likely influence the performance of leaf phenology models driven by meteorological data. Therefore, by using PRI or CCI to assess photosynthetic phenology, dependence on indirect meteorological data for modelling photosynthetic phenology is minimized. Not only may PRI or CCI reduce uncertainty, but their potential to drive a model based on remote sensing products may also improve observations for site-specific heterogeneity. Such observations may not correspond well with coarser meteorological data in remote regions. Considering the increasing availability of satellite based CCI, our results hold promise for improving the large-scale detection of photosynthetic phenology.

## Materials and Methods

### Site description

This study was conducted at two long-term carbon monitoring forest stands at the Turkey Point Observatory in Ontario, Canada, which are part of Global Water Futures Program and Global Fluxnet. The evergreen needleleaf forest (ENF; CA-TP4; 42°42’36.6”N, 80°21’26.6”W) is a homogenous eastern white pine (*Pinus strobus* L.) stand whose trees were planted in 1939. The deciduous broadleaf forest stand (DBF; CA-TPD; 42°38’07.2”N, 80°33’27.8”W) is a >90-year-old naturally regenerated but managed forest. It is dominated by white oak (*Quercus alba* L.) and red maple (*Acer. rubrum* L.), which make up over 50% of the forest, and eastern white pine (5%); the remaining species consist of American beech (*Fagus grandifolia* Ehrh.), black (*Q. velutina* Lam.) and red oak (*Q. rubra* L.), and white ash (*Fraxinus americana* L.). For both forests, the soil type is classified as Brunisolic Gray Brown Luvisol in the Canadian System of Soil Classification ^46^, which is defined as a very fine sandy loam with low moisture holding capacity. Further site and flux and meteorological observations details are given in Table S2 and in Arain and Restrepo-Coupe ^47^ and Beamesderfer, et al. ^48^.

### Eddy covariance flux station

Flux and meteorological data measurements were made at the top of scaffolding towers from 2003 at ENF and 2012 at DBF using Eddy Covariance (EC) systems. Both EC systems used a closed-path gas analyzer (LI-7000 at the ENF and LI-7200 at the DBF site, LI-COR, Lincoln, NE, USA) and a three-dimensional sonic anemometer (CSAT3, Campbell Scientific, Edmonton, AB, Canada). Meteorological data included temperature and relative humidity (HMP45C, Campbell Scientific Inc.); four components of radiation (CNR1 and CNR4 Net radiometers, Campbell Scientific Inc.); soil moisture (CS650, Campbell Scientific Inc.); and precipitation (Geonor T200B at the ENF site and heated CS700 at the DBF site, Campbell Scientific Inc.). Flux data were filtered, de-spiked, gap-filled and processed as described in Arain and Restrepo-Coupe ^47^ following standard algorithms to provide half-hourly values of GPP (GPP_obs_) and meteorological data.

Downward PAR (PAR_dn_) and reflected PAR (PAR_rfl_) were measured at the top of the flux towers at the ENF and DBF sites (LI-200S, LI-COR Inc. at ENF and PQS1, Kipp and Zonen B.V. at DBF). Below canopy downward PAR was also measured at 2m height at both sites to capture incoming radiation (PAR_dn_) and the other facing downwards to capture outgoing radiation (PAR_rfl_). A below-canopy sensor was located 2 m above the ground facing upwards to capture PAR transmitted through the canopy to the ground (PAR_t_). There was no below canopy downwards facing PAR sensor for soil reflectance, so we assumed soil to be black with little light reflectance, since our forest stands were closed during the growing season. Using half-hourly PAR measurments, *f*_APAR_ was calculated as:

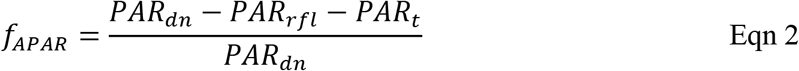

### Measurements of spectral reflectance

Canopy-scale NDVI, NIR_v_, PRI and CCI were estimated using NDVI and PRI spectral reflectance sensors (SRS, Meter Group Inc., Pullman, WA, USA) that were mounted at the tops of the flux towers. Reflectance was measured every minute, and 5-minute averages were recorded. Each site was equipped with three target NDVI and PRI sensors with a field of view of 20°, and two reference NDVI and PRI sensors with a field of view of 180°, to account for incoming radiation. The target sensors were set up facing downwards at a 45° angle facing east, south and west from the flux towers. The reference sensors were mounted near the target sensors so they were unaffected by any shading. The NDVI and PRI sensors measured spectral reflectance at 630 and 800 nm, and 532 and 570 nm, respectively. Each waveband was normalized by dividing the target values by the average of the reference values from the two reference sensors. Sensor cross-calibration was performed to further correct the data, using a Teflon white reference calibration panel that was placed under the target sensors according to Gamon, et al. ^49^. Calibration points were recorded for a minimum of 15 minutes. This procedure was repeated every 2 to 3 months in 2015 and every 3 to 4 months for three years starting in 2016. Using these corrected values, NDVI and PRI were derived from the respective sensors; NIR_v_ was derived from the NDVI sensors; and CCI was derived from a combination of wavebands from the NDVI and PRI sensors, which were calculated as:

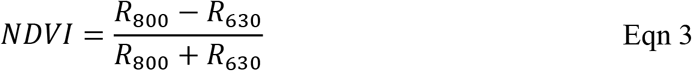

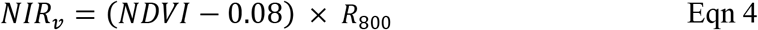

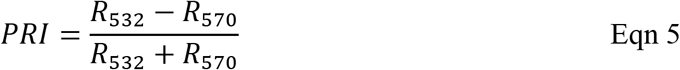

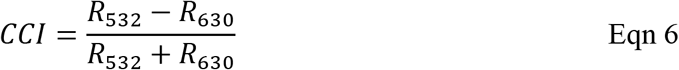

where R indicates the waveband in nm. The recorded 5-minute data points were further averaged to 30-minute aggregates to match flux data.

### Estimating GPP using a LUE model based on NDVI and meteorological data

The LUE model from Yuan, et al. ^21^ is based on the use of remotely sensed NDVI and meteorological data obtained from the flux tower (LUE_Met_) described in Eqn 7 below and in Table S3. *f*_APAR_ was calculated according to Eqn 2 using data from a 5-hour window near solar noon from 11:00 to 16:00 hours to ensure maximum light levels.

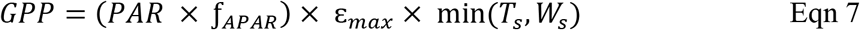

Realized ε was calculated using an invariant maximum ε (ε_max_, g C m^−2^ MJ^−1^ absorbed PAR) shared across sites and biomes reduced by non-optimal air temperature (T_s_) or water stress (W_s_). We used 2.14 g C m^−2^ MJ^−1^ absorbed PAR as εmax from Yuan, et al. ^21^. The non-optimal air temperature factor, T_s_, was estimated using minimum (T_min_), maximum (T_max_) and optimum air temperature (T_opt_), which were set as 0, 40 and 36°C, respectively, and observed air temperature (T) from the meteorological measurments on top of the flux towers (Eqn 8). The canopy water stress factor, W_s_, was estimated using latent heat flux (LE) and sensible heat flux (H) from EC (Eqn 9). The contribution to the ε_max_ reduction factor by non-optimal air temperature or water stress was based on the assumption that ε_max_ would be affected only by the most limiting factor at any given time based on Liebig’s law ^21^. Therefore, only the most limiting contribution of T_s_ or W_s_ was used (i.e., the smallest value indicating a stronger negative impact).

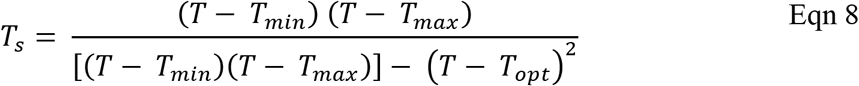

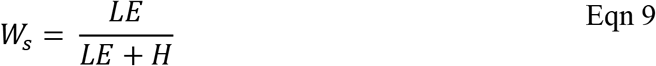

### Estimating GPP using LUE models based on PRI and CCI

We used several vegetation index-based LUE models (LUE_NDVI x PRI_, LUE_NDVI x CCI_, LUE_NIRv x PRI_, and LUE_NIRv x CCI_, see Table S3 for model details). All vegetation index-based LUE models used NDVI derived from SRS data as a proxy for *f*_APAR_ or NIR_v_ as a proxy for APAR with PRI or CCI as proxies for ε (Fig. S6). Based on Eqn 1, we multiplied PAR from the flux tower with NDVI as a proxy for *f*_APAR_ or used NIR_v_ as a direct proxy for APAR and multiplied a normalized (0-1) PRI or CCI as a proxy for ε to estimate an uncalibrated GPP (unitless). The uncalibrated GPP was used to determine a linear regression equation with GPP_obs_ to calibrate GPP from the vegetation index-based LUE models. Prior to normalization, PRI and CCI were filtered using minimum and maximum thresholds for PRI (−0.5 to 0.5) and CCI (−0.5 to 0.5). Additional data filtering included limiting PRI and CCI to PAR values greater than 100 μmol m^−2^ s^−1^. Data during rainfall events were also removed, as rain confounds spectral reflectance signatures detected by the SRS sensors. Last, GPP estimates were automatically set to 0 μmol m^−2^ s^−1^ at temperatures below 0 °C.

### Estimating GPP using a MODIS LUE model

We used the 8-day 500 m GPP product (MOD17A2H) from the MODIS collection 6 land products (MODIS_GPP_) ^50^. MODIS_GPP_ is based on the LUE model using satellite-based NDVI as a proxy for *f*_APAR_ and biome specific limitations of ε from meteorologically based temperature and vapor pressure deficits (VPD) ^16^. We extracted the 500 m pixel matching the coordinates of our sites. We chose to use only the 500 m pixel to cover the footprint of the flux towers. Surrounding areas contained mixtures of other forest types and agricultural land, which would not represent the EC footprint.

A MODIS CCI-based LUE model was used at a 1-km^2^ spatial resolution. Corrected MODIS multi-angle implementation of atmospheric correction algorithm (MAIAC) surface reflectance based on bidirectional reflectance distribution function (BRDF) ^51^ was used to calculate NDVI, NIR_v_ and CCI, which were calculated as:

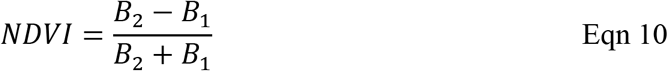

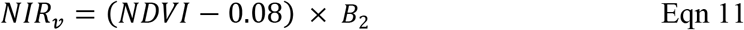

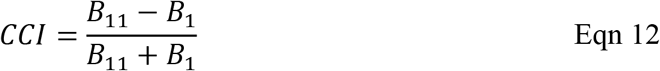

MODIS vegetation index-based LUE models (LUE_NDVI x CCI_ and LUE_NIRv x CCI_), based on Eqn 1, used NDVI as a proxy for *f*_APAR_ with flux tower PAR or NIR_v_ as a proxy for APAR multiplied by CCI as a proxy for ε. MODIS reflectance was screened for cloud-free days. The details of all MODIS LUE are provided in Table S3.

### Estimating GPP using the process-based model JULES

We used the land surface model JULES v4.4 to simulate GPP at the canopy-scale. JULES is a process-based model which incorporates representations of energy balance, radiation, hydrology, soil and vegetation processes and their interactions ^52,53^. Vegetation and soil parameters were prescribed using site-specific information (Table S2). A phenology model that represents the behavior of cold-deciduous vegetation based on a chilling days parameterization ^52,54^ was used for the DBF site. JULES was driven using half-hourly meteorological data obtained from the flux tower at each site. We used air pressure (Pa), specific humidity (kg kg^−1^), air temperature (K), downward shortwave radiation (W m^−2^), downward longwave radiation (W m^−2^), precipitation (kg m^−2^ s^−1^), and wind speed (m s^−1^) as the driving variables. Two variations of JULES were used and are described in detail in Table S3: JULES-Standard (JULES_Std_), which estimates photosynthesis based on Collatz, et al. ^55,56^, and a modified JULES (JULES_Farq_), which uses the Farquhar photosynthesis model ^6^. For JULES_Farq_, we used leaf-scale observations to parameterize the temperature response of maximum rate of Rubisco-mediated carboxylation (*V*_cmax_) and the maximum rate of carboxylation limited by electron transport (*J*_max_) ^57^

### Plant physiological measurements used for JULES model parameterization

We used a LI-6400XT portable photosynthesis system (LI-COR, Lincoln, NE, USA) to conduct A/Ci curves to estimate *V*_cmax_ and *J*_max_ for JULES_Farq_ parameterization. A/Ci curves were conducted every 2-3 weeks during the spring and autumn, and every 4-5 weeks during the summer and winter from March 2015 to the end of 2016 using canopy access towers at both ENF and DBF. Assimilation was assessed after 10 minutes of exposure to CO_2_ levels of 400, 200, 100, 50, 400, 400, 800, 1200 μmol mol^−1^ CO_2_. Measurements were taken at light levels of 1500 μmol quanta m^−2^ s^−1^, relative humidity 55% to 65% and ambient temperatures. *V*_cmax_ and *J*_max_ were estimated based on the C_3_ photosynthesis equations by Farquhar, et al. ^6^. All *V*_cmax_ and *J*_max_ data were pooled together to account for the temperature dependence in the JULES_Farq_ model ^57^; see Table S2 for temperature response parameters. Three trees of eastern white pine (for the ENF) and red maple and white oak (for the DBF) were measured to represent the dominant species of each forest stand.

Leaf nitrogen content was used as an input in both JULES_Std_ and JULES_Farq_. Leaf samples were collected and dried at 80°C for 2 days. Dry leaf samples were homogenized using a grinding mill and packed into tin capsules for nitrogen analysis (ECS 4010 CHNSO ANALYZER, Costech Analytical Technologies Inc., Valencia, CA, USA). From the same trees selected for the A/Ci curves, three leaves from the top of the canopy were collected approximately every 3 weeks from June to August in 2015 and 2016.

### Estimating daily GPP and phenological dates

For GPP_obs_ and all models, we determined daily GPP using sums from a noontime period of 11h to 16h. This was to limit the influence of sun angle on vegetation indices ^58^. We estimated the dates of the start of growing season (SOS) and end (EOS), and the start of the peak of the growing season (SOP) and end (EOP) for each model, and compared these against the phenology dates derived from GPP_obs_. Evaluation of SOS and EOS is based on the difference between observed and modelled dates and denoted as Δ SOS and Δ EOS. Phenological dates were estimated using a fitted seven-parameter logistic function according to Gonsamo, et al. ^36^. Each season was defined according to the estimated phenological dates of GPP_obs_. Spring was defined as the period between SOS and SOP, summer as the period between SOP to EOP, autumn as the period between EOP to EOS, and winter as the period between EOS and SOS. The length of the season was defined as the period between SOS to EOS. More details are given in Beamesderfer, et al. ^59^.

### Statistics

The coefficient of determination (R^2^), bias, root mean square error (RMSE), mean absolute error (MAE), Akaike Information Criterion (AIC) and Bayesian Information Criterion (BIC) for each GPP model were analyzed in RStudio (Version 1.0.136).

## Acknowledgments

This work was supported by funding to IE from NSERC (Grant RGPIN-2015-06514), CFI (Grant 27330), and Ontario Ministry of Research and Innovation (Grant ER10-07-015); funding to MAA for flux measurements was provided by NSERC Discovery, Global Water Futures program and the Ontario Ministry of Environment, Conservation and Parks; LMM acknowledges funding from UK Natural Environment Research council through the UK Earth System Modelling project (NE/N017951/1) and the Montane-Acclim project (NE/R001928/1); CYSW acknowledges The Ann Oaks Doctoral Scholarship awarded by the Canadian Society of Plant Biologists and the Graduate Student Research Award from the University of Toronto Centre for Global Change Science. Support from Long Point Regional Conservations Authority for the CA-TPD site and the Ontario Ministry of Conservations and Forestry as well as St Williams Conservation Reserve Community Council for the CA-TP4 site are also acknowledged. We would like to thank Yazad Bhathena for assistance in field data collection; Petra D’Odorico for help with the phenology fitted double logistics function; Felix Chan, Eric Beamesderfer and Myroslava Khomik for support and advice at Turkey Point Observatory; and Steve Garrity, METER Group and SpecNet for providing the SRS sensors used throughout the study. LMM was supported by the UK Natural Environment Research Council (NERC) research grants NE/R001928/1 and NE/N017951/1. The authors thank Emily Wheeler, Boston, for editorial assistance.

## Author Contribution

CYSW and IE developed the initial framework and objectives of the study. CYSW lead the field data collection and analysis. Eddy covariance flux, meteorological and biometric data for both sites were contributed by MAA. JULES parameterisation and simulations were performed by CYSW with support from LMM. All authors contributed to the writing of the manuscript.

